# Multi-Drug Artificial Neural Network Models for Predicting HIV-1 RTI Resistance

**DOI:** 10.1101/2024.05.03.592325

**Authors:** Sumeyye Yilmaz, Huseyin Tunc, Murat Sari

## Abstract

Drug resistance is a major barrier to effectively treating HIV/AIDS, necessitating the exploration of novel drugs and an understanding of resistance mechanisms. Genotypic and phenotypic tests, while common, are costly and time-consuming. To address these challenges, machine learning models that effectively generalize available data have been leveraged. This study aims to create multi-drug artificial neural network (MD-ANN) models based on drug-isolate-fold change (DIF) to predict the drug resistance profiles of HIV-1 reverse transcriptase inhibitors (RTIs) using the Stanford HIV Drug Resistance Database. Unlike existing isolate-fold change (IF) models, the DIF-based models can test novel RTIs against a given mutant strain. It has been shown that the DIF-based models have competitive predictive capabilities in single-drug benchmarks compared to IF-based models while taking advantage of molecular learning to test novel RTIs. The DIF-based models can rank ten FDA-approved inhibitor pairs according to their drug resistance scores. We validate our observations with an external test set consisting of drug resistance scores for novel RTIs in the CheMBL database. By combining the Stanford dataset and the external dataset, our DIF model achieves impressive classification and regression results with an accuracy of 0.774, AUC of 0.859, and **r** of 0.701 on the test set. To demonstrate how our DIF models learn from drug molecules, we construct a null model that only takes into account isolate representations. This null model yields an accuracy of 0.641, AUC of 0.701, and **r** of 0.470. We highlight the significant contribution of drug information in improving the predictive accuracy and reliability of the DIF model. The DIF models have the potential to facilitate the testing of new inhibitors across various isolates.

## 1 Introduction

Human Immunodeficiency Virus (HIV) is an infectious and chronic disease that has serious effects on the immune system. It infects and destroys immune system cells, which are the body’s natural defense mechanism, and white blood cells such as CD4+ T cells [1]. Initially, HIV-positive individuals may remain unaware of their infection due to the absence of early symptoms associated with HIV. However, as the virus spreads throughout the body and the CD4+ T cell count decreases over time, the immune system weakens, making the person more vulnerable to recurrent opportunistic infections, cancers, and severe health problems. As the HIV infection advances, it leads to the development of Acquired Immunodeficiency Syndrome (AIDS). According to the World Health Organization (WHO), 39 million people are living with HIV worldwide in 2022 and 1.3 million new cases have been reported [2]. Antiretroviral drugs are used for treatment in 76% of HIV-positive patients [2].

By employing antiretroviral therapy (ART), viral replication is suppressed to undetectable levels, resulting in a deceleration of viral dissemination within the organism [3, 4]. The use of ART augments both the lifespan and the overall well-being of individuals [5–7]. However, ART cannot completely eliminate the viral load, and therefore there is no definitive treatment for HIV at the moment [8]. Since HIV is a chronic disease, its treatment requires regular drug use throughout life. Drug resistance is a primary reason for ART failure. It occurs due to increased drug use in the body, the high mutation rate of HIV, suboptimal drug use, poor adherence, or spatial heterogeneity [9–17]. As drug resistance poses a significant obstacle to effective HIV/AIDS treatment, it is crucial to explore new drugs and gain a comprehensive understanding of resistance mechanisms.

Antiretroviral drugs, particularly nucleoside reverse transcriptase inhibitors (NRTIs) and non-nucleoside reverse transcriptase inhibitors (NNRTIs), play a crucial role in combating HIV [7, 18, 19]. NRTIs and NNRTIs specifically target the reverse transcriptase enzyme, which represents the initial and crucial stage of the virus’s life cycle [20, 21]. NRTIs function by mimicking natural nucleosides, participating in DNA synthesis, and subsequently halting DNA formation. Conversely, NNRTIs interact with the reverse transcriptase enzyme, modifying its structure and rendering the enzyme inactive, thereby preventing the generation of DNA [20, 22]. This dual mechanism of action exhibited by NRTIs and NNRTIs underscores their paramount significance in curbing HIV replication. Contemporary HIV treatment protocols consider combinations comprising at least two different NRTIs and at least one NNRTI as indispensable components.

RTI drug resistance can be directly assessed using time-consuming and costly genotypic testing [23]. Predictive mathematical models provide quick and reliable solutions in understanding the genotype-phenotype relations [6, 22, 24**?** –27]. Various machine learning models such as decision trees [25], linear regression [28], support vector machines [29], support vector regressions [22, 30], k-nearest neighbors [24], artificial neural networks [6, 26, 31**?**], and random forest algorithms [24, 31] have been proposed to predict mutational effects in drug resistance to HIV inhibitors. Additionally, various genotypic interpretation algorithms such as Stanford HIVdb [27, 32], HIV grade [33], REGA [34], and ANRS [35] have been derived to tackle the same problem. The rule-based algorithms necessitate ongoing updates to their existing rules and exhibit reduced accuracy when confronted with intricate mutational patterns within sequences [36]. Existing rule-based and machine-learning algorithms are designed to identify and interpret HIV drug resistance for a specific inhibitor. However, these approaches often require substantial computational resources and can be limited by the complexity of mutational patterns.

In this study, we propose a multi-drug artificial neural network (MD-ANN) framework to predict the drug resistance scores of HIV-1 RTIs, utilizing molecular fingerprints and mutational information (Figure 1). Data from RTIs in the Stanford HIV Drug Resistance Database are employed as both training and test datasets for model construction. Unlike existing isolate-fold change (IF) models, DIF models incorporate both inhibitor descriptors and mutational genotype information, enabling predictions of fold changes across any isolate and RTI, including newly discovered inhibitors. These DIF-based models have shown competitive predictive capacity in single-drug benchmarks compared to IF-based models by leveraging molecular learning. They can also rank ten FDA-approved inhibitor pairs by their drug resistance scores. Significantly, our DIF models effectively classify strains as resistant or susceptible, a finding supported by external validation with a test set of drug resistance scores for various RTIs.

**Fig. 1.**
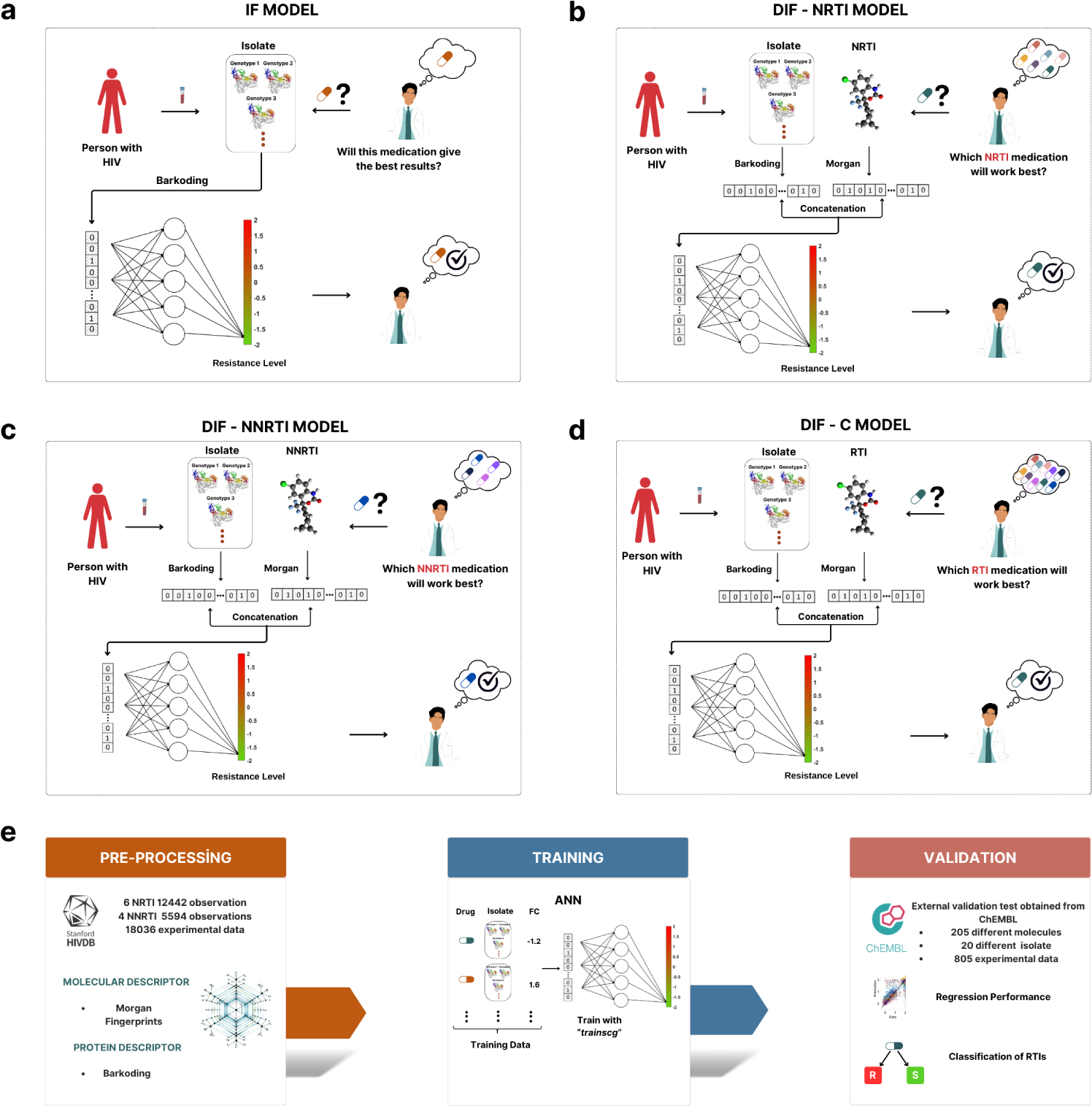
Overview of the study. a) The IF models use isolate information obtained from the patients to predict how this isolate results in resistance to a specific inhibitor. This methodology is well-established in existing literature, as exemplified by references [6, 22, 24, 25, 27, 37]. b) DIF-NRTI models combine information about the isolate and NRTI drugs to forecast treatment outcomes, focusing on specific changes in the virus and drug properties. This model predicts which medication might work best by examining detailed information on the isolate and chemical characteristics of the drugs. c) DIF-NNRTI models are similar to the model in (b), the purpose of this model is to predict the efficacy of drugs; however, it is specifically designed for the NNRTI class of medications. d) DIF-C models are a comprehensive and versatile model designed to evaluate the effectiveness of all types of reverse transcriptase inhibitors (NRTIs and NNRTIs). It uses a detailed profile of viral mutations and RTIs to predict the resistance score of a given RTI. e) The process of model building involves several key stages, starting with the preparation of data, followed by training the neural network, and culminating in the validation of predictions against real-world outcomes. This sequence of steps ensures that the models are both reliable and accurate for predicting HIV treatment outcomes.

## 2 Materials and Methods

### 2.1 Data Descriptions

Filtered genotype-phenotype data for six NRTIs and four NNRTIs are obtained from the Stanford HIV Drug Resistance Database (Figure 1) [38]. The data analysis involved evaluating mutations observed in response to NRTIs and NNRTIs, along with their corresponding fold-change values. Table S1 provides a summary of this analysis, presenting details such as the number of mutations, maximum and minimum fold-change values, mean and standard deviation values, mode value, and first and third quartile values for each inhibitor. Resistance to RT inhibitors is indicated by an elevated fold-change (FC) in *IC*50 (the required drug concentration to inhibit 50% of virions) values compared to wild-type HIV. This involves dividing mutant *IC*50 values by wild-type *IC*50 values, providing a clear measure of the level of resistance exhibited by the mutants [24, 30].

In the analysis conducted, a total of 1352 distinct NRTI mutations and 1365 distinct NNRTI mutations were identified. Since both drug groups target the same enzyme, it is observed that they share common mutations. By combining the data from the two inhibitors, a total of 1388 specific mutations are obtained. To provide a clearer understanding of the data, the inhibitor data is separately analyzed based on the number of isolates indicating the presence of a mutation. The distribution of these mutations within each inhibitor and some statistical values are discussed in Figure 2 and Table S1.

**Fig. 2.**
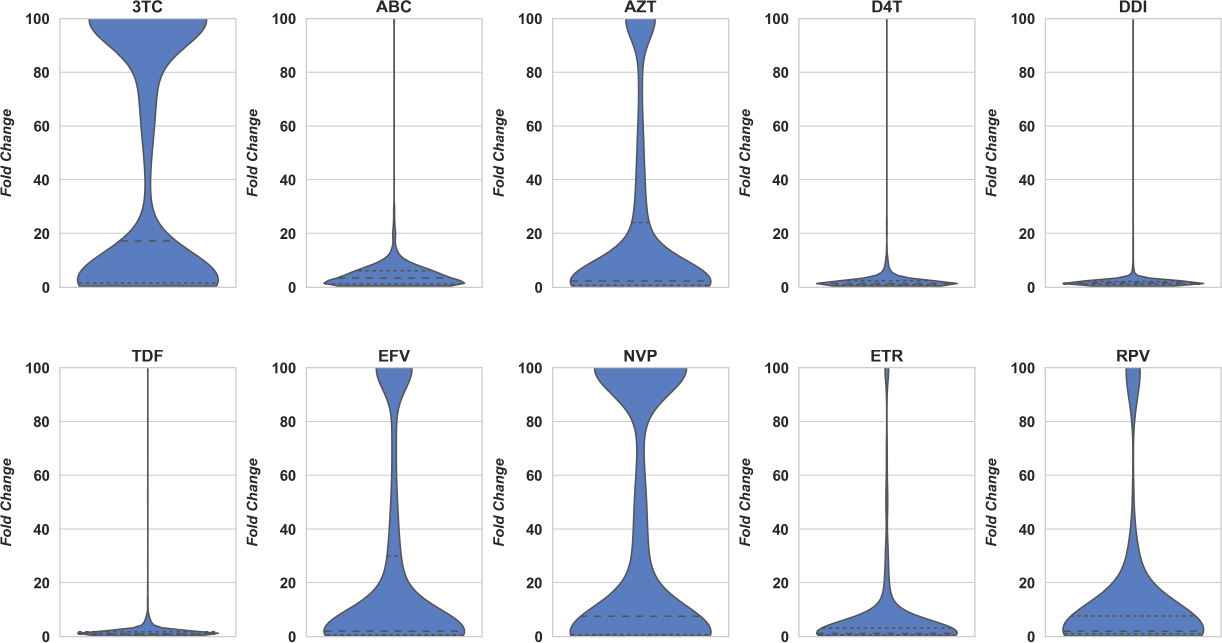
Distribution of fold change values obtained for each inhibitor.

Among the 1388 unique mutations obtained from the inhibitors, we conducted an analysis to determine the frequencies of these mutations within the dataset. From all the mutations analyzed, we identified the top 30 mutations with the highest occurrence rates across the entire dataset. These mutations represent the most commonly observed variations within our dataset. In Figure S1, we visually represent these mutations. Furthermore, in Figure S1, we categorize these mutations based on the residues they affect. Among the numerous mutations, three specific variants, namely R211K, K122E, and M184V, are the most prevalent in occurrence frequency [39, 40].

### 2.2 Representation of Isolates

In the dataset comprising six NRTIs, we observed a total of 1352 distinct mutations. Similarly, in the dataset involving four NNRTIs, we identified 1365 unique mutations. When both drug classes are evaluated together, a total of 1388 unique mutations are documented due to the presence of shared mutations between the two drug classes. In this study, we employed the widely used binary barcoding technique from the literature to represent isolates present in genotype-phenotype datasets of various HIV-1 inhibitors [26, 28]. For this reason, 1352, 1365, and 1388-dimensional representation vectors are created using 0s and 1s depending on the existence of mutations in an isolate. Suppose that 1352, 1365, and 1388 unique mutations produce vectors *X* = [*x*_1_, *x*_2_,…, *x*_1352_], *Y* = [*y*_1_, *y*_2_,…, *y*_1365_] and *Z* = [*z*_1_, *z*_2_,…, *z*_1388_], respectively. Here, *x_i_, y_i_*, and *z_i_* are elements of mutation sets occurring in the dataset. For example, *x*_1_ indicates the occurrence of the A114V mutation, and *x*_1301_ indicates the occurrence of the V90A mutation. In this case, an isolate *I_j_* = *{A*114*V, V* 90*A}* can be barcoded as *X* = [1, 0,…, 0, 1, 0,…, 0]; where only the first and one thousand three hundred first positions take a value of one and the remaining entries have a value of zero. Any isolate *I_j_* = *{a*_1_, *a*_2_,…, *a_n_}* can be represented with *X_j_*= *{x*_1_, *x*_2_,…, *x*_1352_*}* where

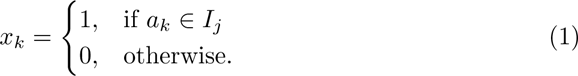

In this way, the isolates undergo conversion into distinctive input vectors for machine learning, consisting of dimensions 1352, 1365, and 1388. Since each mutation occupies a distinct location in the entry vectors created, the binary barcoding method offers the advantage of encoding multiple mutational alterations in amino acids. For instance, let us consider the mutations V106I and V106L represented by *x*_1205_ and *x*_1206_, respectively. In this case, the isolate *I* = *{V* 106*I, V* 106*L}* can be represented as *X* = [0, 0,…, 0, 1, 1, 0,…, 0]. It is important to note that the genotype fold change measurements are made on the virus population, so it is possible to find multiple mutations for a single residue.

### 2.3 Representation of Inhibitors

We generate molecular representations of RTIs using Morgan fingerprints that effectively characterize the molecular structures [16]. Morgan fingerprints are favored in machine learning models for their efficient molecular vector representation. The RDKit [41] environment of the Python program converts the SMILES representations of inhibitors into binary vectors, with variations in bit and radius values. Vectors with different radii and bits are dimensionally reduced to highlight unique characteristics. Figure S2 illustrates the original and reduced dimensions under different settings. The mean *r* values, obtained using Morgan fingerprints represented by various bits and radii with the DIF and DIF-C models, are displayed in Figure S3. The selection of representation is informed by these observations.

### 2.4 Neural Network Construction

We introduce four distinct neural network models, primarily differentiated by the learning mechanism based on the isolate and inhibitor representations. The DIF foundational model functions as a supervised neural network, designed to analyze the resistance level of a reverse transcriptase isolate when exposed to a reverse transcriptase inhibitor (RTI). DIF model has three derivatives DIF-NRTI (Figure 1b), DIF-NNRTI (Figure 1c), and DIF-C (Figure 1d). On the other hand, IF models are designed for a specific RTI (6 IF models for NRTIs and 4 IF models for NNRTIs) by mapping isolate information to resistance score (see Figure 1a).

Over the years, IF models have been developed to predict changes in HIV-1 phenotypes based on genotype information specific to each HIV-1 inhibitor [6, 22, 24, 25, 27, 37]. Leveraging the extensive genotype-phenotype data for various HIV-1 inhibitors available in the Stanford HIV Drug Resistance Database, these models have been successfully constructed using a range of machine learning and deep learning algorithms. The effectiveness of these models varies depending on the dataset size and the nature of the experimental data. For instance, the 3TC inhibitor is modeled with a high degree of accuracy (*R*^2^ = 0.975), whereas the model for the ABC inhibitor is less accurate (*R*^2^ = 0.614) [42]. This raises an important question: Can a machine learning model be constructed to predict the drug resistance score for any inhibitor within an inhibitor group even with limited number of available inhibitors? Tunc *et al.* [26] explored this question in the context of protease inhibitors, demonstrating that DIF models can predict resistance profiles for novel PIs. In this study, we aim to systematically expand this approach and develop DIF models for reverse transcriptase inhibitors.

The DIF-C model is distinguished by its utilization of drug-isolate-fold change data across all reverse transcriptase inhibitors (RTIs). It incorporates a comprehensive 1594-dimensional input, with 1388 dimensions representing isolates and 206 dimensions for Morgan fingerprints. This model is capable of predicting the resistance profile of an RTI for a specific isolate. The DIF-NRTI and DIF-NNRTI models are adapted for the NRTI and NNRTI group of inhibitors. The DIF-NRTI model features an input dimension of 1478, comprising 1352 dimensions for isolates and 126 for Morgan fingerprints. Similarly, the DIF-NNRTI model has an input dimension of 1480, with 1365 dimensions dedicated to isolates and 115 to Morgan fingerprints. These models, as specialized versions of the DIF-C model, are focused exclusively on their respective RTI groups. Additionally, the IF-NRTI and IF-NNRTI models, specifically designed for each inhibitor type, have input dimensions of 1352 and 1365, respectively, corresponding to mutation-specific data.

The DIF and IF models are developed in MATLAB (version 2022a) with GPU acceleration and using the Machine Learning and Deep Learning toolbox, operate on isolate-inhibitor input (isolate input for IF models) and logarithmic fold-change output. The ANN architecture comprises a single hidden layer with five neurons and a single output, tuned through hyperparameter optimization (see Figure S4). The models employ hyperbolic tangent-sigmoid and linear activation functions. The dataset is partitioned into training, testing, and validation sets at ratios of 80%, 10%, and 10%, respectively. During the training phase, the models aim to minimize the mean squared error (MSE) between the predicted values and the actual numerical observations. This optimization is achieved through the *trainscg* optimization built-in function of the MATLAB programming environment. Each ANN model undergoes a 10-fold cross-validation process to ensure the statistical robustness of its performance metrics. To enhance the generalizability of the ANN models, we implement a 10 *×* 5 ensemble learning strategy. In this approach, we train ten models and select the one with the lowest MSE on its internal test set. Repeating this process five times, we acquire five representative models. The predictions for the validation set are then generated by averaging the predictions from these models.

## 3 Results

### 3.1 Regression Performance

We critically evaluate the regression performance of the generated models to determine their impact on our prediction models for HIV drug resistance. For clinical applications, particularly in assessing resistance to specific reverse transcriptase inhibitors (RTIs), IF models may be preferable due to their targeted design. However, when it comes to exploring novel reverse transcriptase inhibitors, the DIF and DIF-C models exhibit distinct advantages. The inherent limitations of IF models preclude their use in analyzing novel inhibitors, a gap that DIF and DIF-C models are specifically designed to address.

The DIF, DIF-C, and IF models are assessed using a 10-fold cross-validation approach. The regression performances of the models are presented using the *r* values (Pearson correlation coefficients), as illustrated in Figure 3. The obtained results demonstrate that our models have accurately learned molecular information from Morgan fingerprints.

**Fig. 3.**
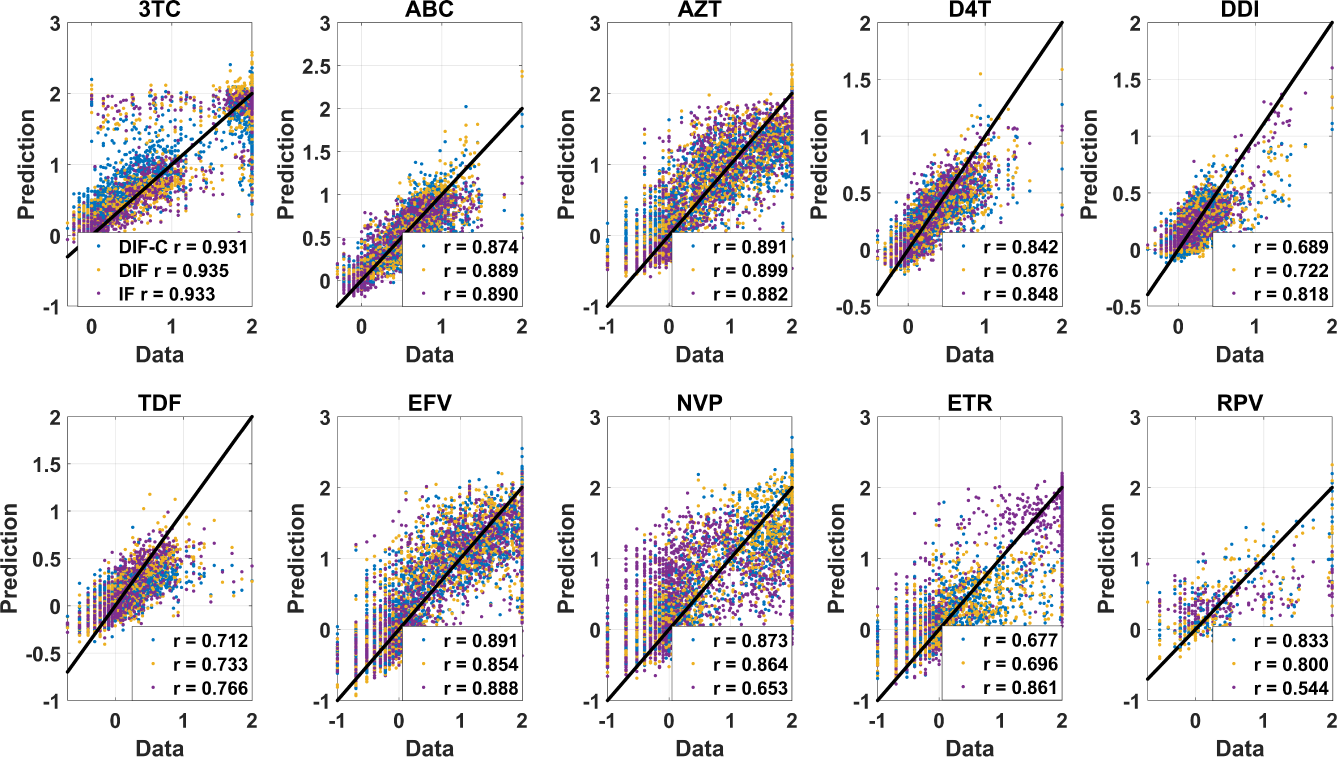
The fold change predictions of IF, DIF-C, and DIF models are shown against actual values for both NRTIs and NNRTIs. The blue mark corresponds to the DIF-C models, the yellow mark corresponds to the DIF models, and the purple marks correspond to the IF models. Models are evaluated using a 10-fold cross-validation procedure and *r* performance values are obtained.

Here we make a comparison to show that DIFs can compete with IFs and give good enough regression performance. The comparative assessment of the DIF-based and IF models revealed a comparable level of performance for NRTI-class drugs. However, a distinctive advantage of the DIF-based models is observed in the case of NNRTI class drugs, with RPV and NVP showing positive responses, contrasted by a lesser effectiveness for ETR. When examining the average *r* performance for NNRTI drugs, it becomes evident that DIF-based models have achieved higher *r* values. These findings indicate that DIF-based models can better adapt to the specific characteristics of NNRTI drugs compared to IF models and can make more effective predictions. Furthermore, an extensive comparative analysis is executed to elucidate the differences in error metrics and regression performances between the IF, DIF, and DIF-C models, as detailed in Table S2. The comparison between the DIF and IF models is particularly emphasized, with *r* and MSE values serving as the primary metrics. The results of this comparative evaluation are graphically depicted in Figure S5, providing a clear visual representation of the performance metrics.

DIF-C models contribute significantly to EFV and NVP inhibitors only from the NNRTI class compared to DIF models, but not to ETR and RPV inhibitors of the same class. This discrepancy can be attributed to the distinct structural characteristics exhibited by the ETR and RPV inhibitors, setting them apart from other molecules. To visually emphasize this distinction, principal component analysis (PCA) serves as a powerful tool, allowing for the visualization of molecules in a two-dimensional space. It enables a clear distinction between ETR and RPV inhibitors and other molecules due to their occupation of different regions. Figure S6 provides a comprehensive understanding of the spatial distribution and separation of these inhibitors from other molecules. Through this visualization technique, the unique structural characteristics and inherent dissimilarity of the ETR and RPV inhibitors are further highlighted, thus bolstering the understanding of their distinct nature and justifying the limited impact of the DIF-C models on these particular inhibitors.

### 3.2 Drug Resistance Tendencies of RTIs

The comparative analysis of resistance or sensitivity to a specific genotype between any two drugs is crucial in the development of new drug molecules, shaping drug design and therapeutic strategies. Our DIF and DIF-C models are designed for this predictive mechanism. Our models focus on assessing which of two drugs exhibits greater resistance compared to the same genotype. Understanding how different genotypes influence drug efficacy and resistance profiles is essential for optimizing treatment strategies and improving patient outcomes. Through the examination of fold-change values corresponding to specific drug-genotype pairs, valuable insights emerge regarding inhibitor resistance profiles. These tendencies serve as crucial indicators of the relative effectiveness of inhibitors against specific genotypes, furnishing essential information for the development of targeted therapies. This knowledge aids in pinpointing effective treatments tailored to individuals with specific genotypes, thereby enabling precise and personalized approaches.

In both the DIF and DIF-C models, a test set comprising 20% of the data from each inhibitor is set aside, while the remaining 80% of the data is utilized for training the models. This approach ensured that a separate and independent dataset is available to assess the performance and generalization ability of the trained models. The test set, which consisted of the excluded 20% of the data, is then used to evaluate the performance of the models. By utilizing the obtained two-dimensional correlation coefficients, our objective is to assess which drug performed better on specific genotypes compared to the other drugs and to quantify the degree of this superiority. The DIF and DIF-C models are employed for the analysis to determine the extent to which they exhibited superior performance.

Among the pairs in the DIF-C models, the 3TC-DDI pair exhibited the highest 2D correlation coefficient of 0.924 (95% CI [0.793, 0.841]). On the other hand, the DDI-ETR pair exhibited the lowest correlation coefficient of 0.606 (95% CI [0.793, 0.841]). The 2D correlation coefficients for the DIF models are 0.939 and 0.647 for the best 3TC-D4T and worst ETR-RPV drug pairs, respectively (95%CI [0.814, 0.862]). These comparisons in the DIF-C and DIF models are detailed in Figures 4-S7.

**Fig. 4.**
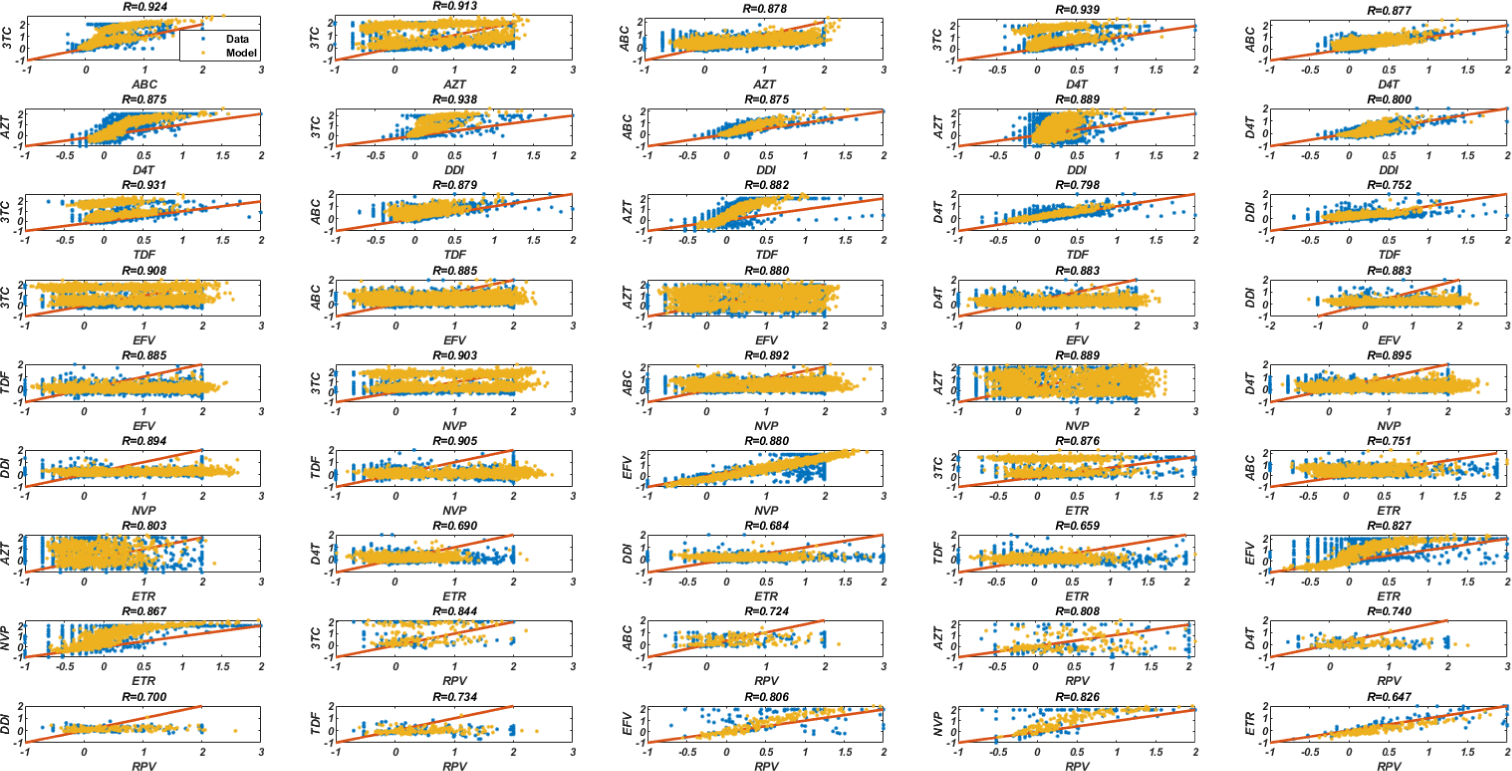
Prediction of fold change tendencies of inhibitor pairs with the DIF models obtained by training NRTIs and NNRTIs separately.

DIF-C models exhibit superior performance compared to DIF models for several drug pairs, such as EFV-NVP and ABC-RPV. The DIF-C models achieve higher 2D correlation coefficients (0.883 and 0.733, respectively) compared to the DIF models (0.880 and 0.724, respectively). This trend continues with other drug pairs like DDI-RPV, TDF-RPV, EFV-RPV, NVP-RPV, and ETR-RPV, where the DIF-C models consistently outperform the DIF models with higher 2D correlation coefficients. These findings indicate that the DIF-C models are more effective in predicting resistance profiles for these specific drug pairs. However, it is important to note that the DIF models demonstrate better performance for the remaining thirty-eight drug pairs, excluding the seven drug pairs where DIF-C models outperform them.

Table 1 allows for a comparative analysis of the average 2D correlation coefficients of the DIF-C and DIF models for different inhibitor classes and all inhibitors. Upon examining the drug resistance tendencies in the table, it becomes apparent that the DIF models have a positive impact on NRTI-NRTI and NRTI-NNRTI combinations, as compared to the DIF-C models. For NNRTI-NNRTI combinations, both the DIF-C and DIF models demonstrate competitive performance. In other words, the DIF-C models contribute positively to drug resistance tendencies prediction for the NNRTI class.

**Table 1.** Mean 2D correlation coefficients according to the NRTI and NNRTI classes predicted by DIF-C and DIF models.

| Drug Classes | Average 2D R |  |
| --- | --- | --- |
|  | DIF-C | DIF |
| **NRTIs-NRTIs** | 0.835 (95% CI [0.799, 0.871]) | **0.877 (95% CI [0.846, 0.907])** |
| **NNRTIs-NNRTIs** | 0.807 (95% CI [0.721, 0.893]) | **0.809 (95% CI [0.721, 0.897])** |
| **NRTIs-NNRTIs** | 0.808 (95% CI [0.770, 0.846]) | **0.822 (95% CI [0.785, 0.858])** |
| **All** | 0.817 (95% CI [0.793, 0.841]) | **0.838 (95% CI [0.814, 0.862])** |

These analyses demonstrate the ability of both DIF and DIF-C models to differentiate resistance potentials among various RTI pairs. While the DIF models generally capture resistance profiles more effectively and provide more accurate predictions, the DIF-C models excel in specific drug combinations. These insights enhance our understanding of the models’ strengths in drug resistance tendencies prediction.

### 3.3 Classification of NRTIs and NNRTIs

DIF-based models are capable of determining the fold change values for each reverse transcriptase inhibitor across various isolates. This prediction task can be formulated as a classification problem, where the goal is to determine the relationship between the fold change values of two potential inhibitors, denoted as *A* and *B*, for a given isolate. Specifically, the classification criterion is defined as, *log*(*FoldChange*[*A, isolate*]) *> log*(*FoldChange*[*B, isolate*]). The resulting relationships between inhibitors and isolates are represented by binary values of 0s and 1s.

To ensure the models’ robustness and prevent overfitting, 5-fold cross-validation techniques are employed during the training phase. This approach allowed for comprehensive training and evaluation of the models across different subsets of the data. The performance of the final model is assessed using the test data, providing an unbiased evaluation of its predictive capabilities. The corresponding receiver operating characteristic (ROC) curves are shown in Figure 5.

**Fig. 5.**
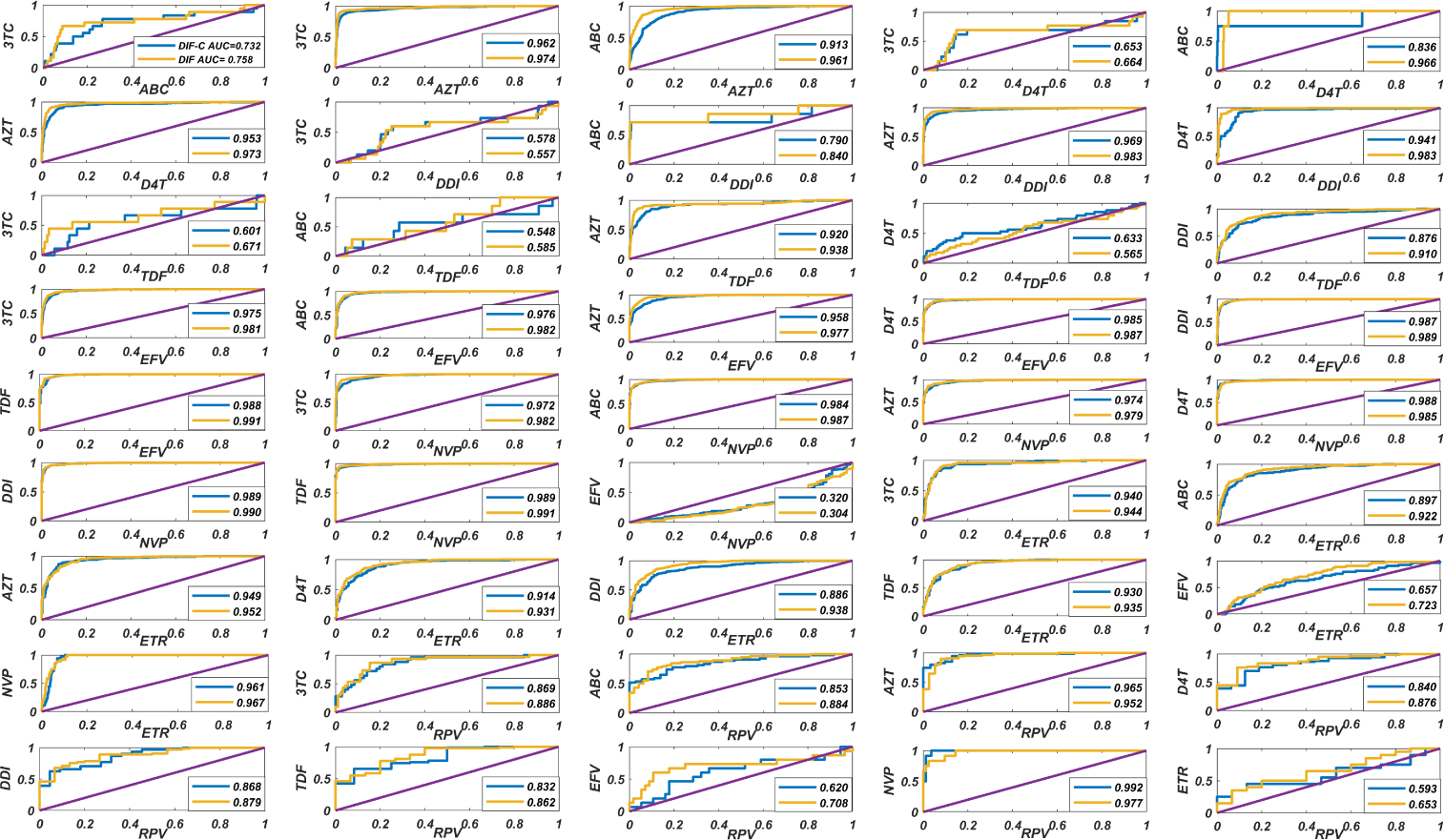
Classification performances of the DIF and DIF-C models are evaluated using the area under the ROC curves (AUC) as a measure of performance. The accuracy of the models is assessed based on the true estimation rate. The DIF-C model’s performance is represented by the blue ROC curve, while the DIF model’s performance is represented by the yellow ROC curve.

Figure 5 displays the values for the area under the ROC curve (AUC) for the DIF-C and DIF models, where NRTI and NNRTI are trained jointly and separately, respectively. The AUC values for the DIF-C models range from 0.992 for the best pair of NVP-RPV to 0.320 (95% CI: [0.809, 0.904]) for the worst pair of EFV-NVP. Conversely, the DIF models yield AUC values of 0.991 for the best pairs TDF-EFV and TDF-NVP. The worst AUC value obtained from the DIF models is 0.304 (95% CI: [0.830, 0.923]) for the EFV-NVP pair. The DIF-C models consistently outperformed the DIF models in terms of AUC across multiple drug pairs. Specifically, the DIF-C models exhibited higher AUC values for the 3TC-DDI, D4T-TDF, EFV-NVP, AZT-RPV, and NVP-RPV drug pairs. This indicates that the DIF-C models are more effective in predicting drug trends for these specific drug pairs. However, it’s important to note that the DIF models showed better AUC performance in the remaining 40 drug pairs excluding 3TC-DDI, D4T-TDF, EFV-NVP, AZT-RPV, and NVP-RPV drug pairs.

Table 2 presents the average AUC values for the DIF and DIF-C models, categorized by drug classes and overall drugs. The DIF models consistently outperformed the DIF-C models, indicating its higher effectiveness in predicting drug resistance within the given dataset. Notably, in the NRTI-NRTI drug class, both models exhibited high AUC values, but the DIF models slightly outperformed the DIF-C models, suggesting its better ability to capture resistance patterns in this specific class. The NRTI-NNRTI drug class showed higher average AUC values for both models, indicating relatively successful prediction of drug resistance in combinations of NRTIs and NNRTIs. Conversely, the NNRTI-NNRTI drug class had lower average AUC values for both models, highlighting the challenges in resistance prediction for this class and the need for further improvements. These findings underscore the varying performance of the DIF and DIF-C models across different drug classes and emphasize the complexities involved in accurately predicting drug resistance. Further analysis and investigation are necessary to gain a comprehensive understanding of the underlying factors contributing to these observed patterns.

**Table 2.**
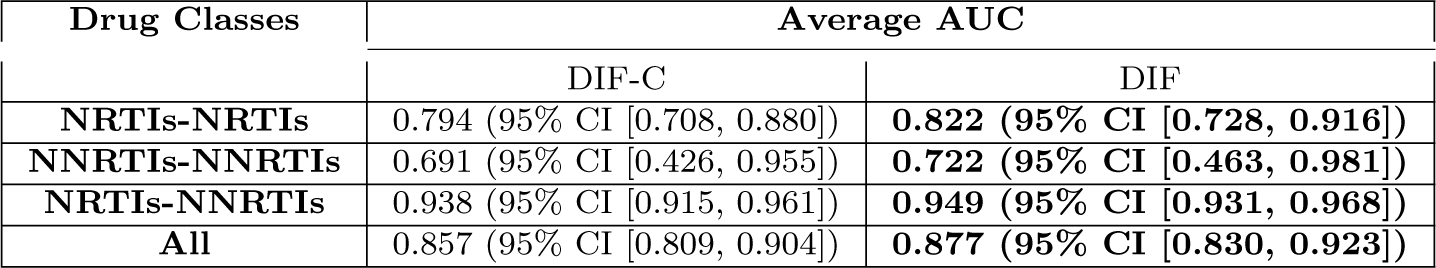
Mean AUC values of DIF and DIF-C models based on inhibitor classes. Performance is evaluated using the area under the ROC curve (AUC) and mean across the values.

| Drug Classes | Average AUC |  |
| --- | --- | --- |
|  | DIF-C | DIF |
| **NRTIs-NRTIs** | 0.794 (95% CI [0.708, 0.880]) | **0.822 (95% CI [0.728, 0.916])** |
| **NNRTIs-NNRTIs** | 0.691 (95% CI [0.426, 0.955]) | **0.722 (95% CI [0.463, 0.981])** |
| **NRTIs-NNRTIs** | 0.938 (95% CI [0.915, 0.961]) | **0.949 (95% CI [0.931, 0.968])** |
| **All** | 0.857 (95% CI [0.809, 0.904]) | **0.877 (95% CI [0.830, 0.923])** |

Table 3 present the mean accuracy, sensitivity, specificity, precision, recall, and F1 score values for drug classes. When examining these tables, it can be observed that the accuracy of the DIF models are higher than the DIF-C models for drug classes. Looking at the average sensitivity and specificity values, the DIF-C models perform better than DIF in the NNRTI-NNRTI drug class. However, in all other cases, the DIF models outperform DIF-C. The DIF-C models show higher average precision and recall values for the NRTI-NRTI and all mean drug classes compared to the DIF models. However, in all other classes, the DIF models have higher average precision and recall values. Analyzing the average F1 score values, it is evident that the NNRTI-NNRTI drug class performs significantly better in the DIF-C models compared to DIF. Overall, it can be concluded that the DIF-C models contribute to the performance measures, particularly in the NNRTI-NNRTI drug class.

**Table 3.** Mean values of accuracy, sensitivity, specificity, precision, recall, and F1 score for the DIF-C and DIF models.

| Metric | Models | NRTIs-NRTIs | NNRTIs-NNRTIs | NRTIs-NNRTIs | All |
| --- | --- | --- | --- | --- | --- |
| Accuracy | DIF-C | 0.928 | 0.814 | 0.883 | 0.889 |
|  | DIF | **0.949** | **0.827** | **0.899** | **0.906** |
| Sensitivity | DIF-C | 0.647 | **0.548** | 0.837 | 0.735 |
|  | DIF | **0.668** | 0.528 | **0.871** | **0.758** |
| Specificity | DIF-C | 0.647 | **0.548** | 0.837 | 0.735 |
|  | DIF | **0.668** | 0.528 | **0.871** | **0.758** |
| Precision | DIF-C | **0.551** | 0.496 | 0.871 | **0.716** |
|  | DIF | 0.459 | **0.521** | **0.890** | 0.704 |
| Recall | DIF-C | **0.551** | 0.496 | 0.871 | **0.716** |
|  | DIF | 0.459 | **0.521** | **0.890** | 0.704 |
| F1 Score | DIF-C | 0.569 | **0.514** | 0.864 | 0.744 |
|  | DIF | **0.734** | 0.375 | **0.889** | **0.775** |

Tables S3 - S4 showcase the evaluation measures utilized to assess the performance of the existing ANN models in capturing the pairwise relationships between the RTI pairs. It can be seen that these models are able to capture the relationships between any pair with high performance. The tables are provided in the order of drugs A and B. The ordering of drugs determines the assignment of positive and negative labels in the classification problem. However, if the order is reversed and drug B precedes drug A, the labels will be assigned differently, potentially leading to a different classification outcome. So, the sensitivity value for drugs A and B corresponds to the specificity value for drugs B and A. The precision value for drugs A and B is equivalent to the recall value for drugs B and A. Consequently, although these values are not explicitly given in the table, they are taken into account when calculating the averages.

### 3.4 External Testing

Testing our models with external datasets serves as a critical indicator of their performance in real-world scenarios. We address the integration of our DIF-C model with real-world data and how this process enhances the overall comprehensiveness of our models. The DIF-C model is trained using both the Stanford dataset and an external dataset. To increase the reliability of the DIF-C model, a 5-fold cross-validation method is applied to the external dataset. In this process, each fold is sequentially used as the test set, with the remaining 4 folds used for training. The performance of the model is evaluated by averaging the results obtained from each fold. This step is of critical importance in demonstrating the practical applicability of our model beyond theoretical limitations.

To create the external dataset, a dataset search was conducted in the ChEMBL [43] database, filtering for compounds with more than 40% similarity to known reverse transcriptase drugs. Throughout the research, molecules with established *IC*50, *EC*50, and *EC*90 measurements in mutants are collected. Notably, filtering for certain inhibitors resulted in finding no molecules at all. From this research, 805 compounds analogous to RTIs such as 3TC, AZT, EFV, NVP, ETR, and RPV are gathered. It is observed that among the 805 inhibitor-genotype-phenotype relations, there were 208 distinct molecules.

Each molecule is associated with a Morgan fingerprint, defined as 512-bit vectors, to generate predictions for our model. We compress these vectors to a more manageable size of 206 dimensions by removing bits that lack informative value. This process has been applied to ten existing drugs, and subsequently, the external dataset was processed through this map. Additionally, we designed a null model by excluding the drug information from the dataset to assess if our DIF-C models learn from inhibitor representations. Our goal was to isolate the models’ learning capacity to genotype-phenotype relationships exclusively. We consider three different model configurations: a null model (Morgan free), Morgan-512, and Morgan-206. The Morgan-512 model uses molecular representations without dimension reduction, whereas the Morgan-206 model is constructed by reducing the dimensions according to the ten RTIs. Given that DIF-C models are more general compared to DIF models, we examined the classification and regression performance of DIF-C models to evaluate their molecular learning capabilities.

The classification of resistance refers to RTIs that are considered resistant when their threshold fold change values are equal to or greater than 3 [24]. The goal of this task is to accurately determine the resistance levels of various molecules—categorizing them as either resistant or susceptible—through the utilization of our models. We evaluate the DIF-C models’ performance in classifying resistance within external datasets by employing several key metrics: accuracy, sensitivity, specificity, precision, recall, F1 score, AUC values, and additionally, the regression metric *r*-value (Table 4). Figure 6 displays the ROC curves for the DIF-C models’ resistance classification performances. This comprehensive assessment underscores our models’ capability to discern molecular resistance levels effectively.

**Fig. 6.**
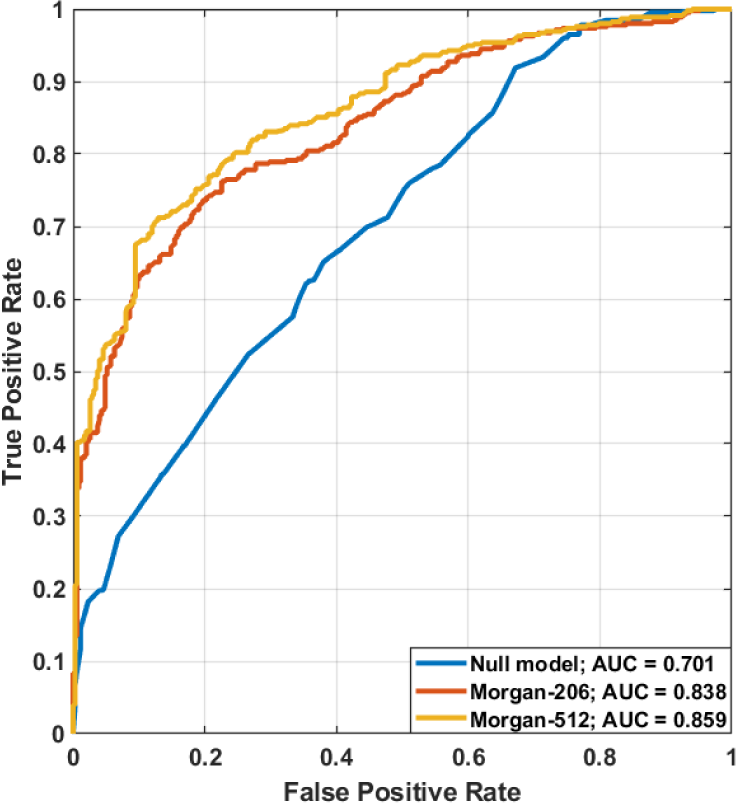
ROC curves and AUC values for the null (Morgan free), Morgan-206, and Morgan-512 model configurations evaluated on the external data.

**Table 4.** Comparison of the DIF-C models (Morgan-512 and Morgan-256) and the IF model (null model) using a five-fold cross-validation approach on the external test data.

| Models | Null model | Morgan-512 | Morgan-206 |
| --- | --- | --- | --- |
| Accuracy | 0.641 | **0.774** | 0.739 |
| Sensitivity | **0.826** | 0.824 | 0.791 |
| Specificity | 0.400 | **0.709** | 0.671 |
| Precision | 0.642 | **0.786** | 0.758 |
| Recall | **0.826** | 0.824 | 0.791 |
| F1 Score | 0.722 | **0.805** | 0.774 |
| AUC | 0.701 | **0.859** | 0.838 |
| $r$ | 0.470 | **0.701** | 0.687 |

The results obtained delineate the performance disparities among the DIF-C model configurations. Notably, the analysis of the null model provides compelling evidence of the critical role played by drug information in the performance of the model. Utilizing drug properties, our model achieved impressive outcomes in resistance classification, with an accuracy of 0.774 and an AUC value of 0.859. Conversely, the exclusion of drug information led to a reduction in accuracy to 0.641 and a decrease in the AUC to 0.701 in the null model, underscoring the significant contribution of drug information to enhancing the predictive accuracy and reliability of the model. The Morgan-512 configuration, across various performance metrics such as accuracy, specificity, precision, F1 score, AUC, and correlation coefficient, consistently displayed superior results. This configuration underscores the superiority of this model configuration in the task of molecular classification, effectively maximizing the capability of the models to accurately classify molecules as resistant or susceptible through the efficient utilization of drug information. On the other hand, despite providing high sensitivity and recall rates, the null model is limited due to its low specificity. This indicates a high false positive rate, which could lead to misleading results in certain instances. The Morgan-206 model, while offering a good balance across performance metrics, does not achieve as high a performance as the Morgan-512. This suggests that the reduction in data dimensionality may adversely affect the learning capability of the models, yet still provides an acceptable level of performance. These findings underscore the importance of drug information in enhancing the performance of the DIF-C model in resistance classification tasks.

## 4 Discussions and Conclusions

We have proposed a multi-drug artificial neural network (MD-ANN) approach based on DIF (Drug-Isolate-Fold Change) for predicting HIV-1 RTI resistance. Training and test data were obtained from the Stanford HIV Drug Resistance Database, following careful preprocessing of the cleaned RTI data. We constructed various models using the NRTI and NNRTI classes. To enhance model reliability and prevent overfitting, techniques such as cross-validation, ensemble learning, and hyperparameter tuning were employed.

This investigation, inspired by the recent study of Tunc *et al.* [26], extends the applicability of DIF models to RTIs. Our study sets itself apart by conducting a more detailed computational analysis, marking a pioneering application of multi-drug ANN models to RTIs. The critical distinction between our developed DIF and DIF-C models and the conventional IF models prevalent in the literature is the incorporation of drug information into predictive models. IF-based models are limited for specific drugs and only learn from genotype representations [6, 24, 25, 28–31, 44]. Our DIF-based models overcome this limitation. Importantly, the aim of this study is not to construct a superior model for existing ten RTIs but to demonstrate that integrating drug molecule information into the DIF model makes predictions for new drug compounds possible. This is achieved by combining isolate and drug structure data in the DIF models, enabling the effective evaluation of new molecules with a given isolate. This attribute enhances the capacity of the DIF-based models to utilize a more diverse dataset, resulting in more comprehensive predictions without compromising the efficiency achieved by IF-based models.

Our research employs a null model approach to critically evaluate the impact of the absence of drug information on the predictive performance of our models. This comparison effectively highlights the significant contribution of drug molecule data to the accuracy and reliability of our predictions. The null model, lacking specific drug information, serves as a baseline to assess the intrinsic value of incorporating drug information into DIF models. Results from this comparative analysis reveal a noticeable decrease in performance metrics such as accuracy, AUC, and correlation coefficient (*r*) when drug information is excluded. Specifically, accuracy, AUC, and r values have been observed to decrease from 0.774, 0.859, and 0.701 in the DIF models to 0.641, 0.701, and 0.470 in the null model, respectively. These findings strongly support the hypothesis that the integration of comprehensive molecular data is crucial for accurately predicting drug resistance profiles.

The DIF and DIF-C models are capable of classifying inhibitors as well as resistant/susceptible strains by leveraging resistance profiles from an external dataset. This method offers a novel perspective by combining two drug groups acting on the same enzyme, thereby facilitating the integration of inhibitory properties into the input side of machine learning models. Such an approach could expedite the identification of effective HIV treatments, conserving time and resources. Notably, for predicting drug resistance scores of FDA-approved NRTIs, IF models are deemed more suitable due to their specific design and the qualified data available for training. However, our findings suggest that for the NNRTIs EFV, ETR, and RPV, the DIF models outperform the IF models, except for the ETR inhibitor within the NNRTI class, where the IF model demonstrates superior accuracy compared to the DIF models.

Our findings provide valuable insights into understanding drug resistance at the molecular level and can contribute to the development of future drug design strategies. The study also suggests that model performance could be further enhanced by employing larger datasets and considering diverse molecular features, thereby enabling the exploration of additional drug classes. This approach could greatly assist in managing drug resistance and developing personalized treatment strategies effectively. The obtained results, bolstered by the use of a null model, highlight the importance of drug information in improving the predictive accuracy and reliability of our models.

## Acknowledgements

The first author would like to thank the Science Fellowships and Grant Programmes Department of TUBITAK (TUBITAK BIDEB) for their support of this academic research.

## Author contribution

Sumeyye Yilmaz: Conceptualization, Methodology, Software, Formal analysis, Investigation, Data Curation, Writing - Original Draft, Writing - Review & Editing.

Huseyin Tunc: Conceptualization, Methodology, Software, Formal analysis, Investigation, Data Curation, Writing - Review & Editing, Supervision.

Murat Sari: Writing - Review & Editing, Supervision

## Data and Software Availability Statement

The codes and data necessary to reproduce the results of our study are publicly available on GitHub: https://github.com/sumeyyeeyilmaz/MD-ANN-hivreversetranscriptase. We downloaded the cleaned HIV reverse-transcriptase genotype-phenotype data from the Stanford drug resistance database: https://hivdb.stanford.edu/. The external dataset was curated using the ChEMBL database: https://www.ebi.ac.uk/chembl/. The GitHub repository includes both datasets, along with codes for preprocessing, training, and postprocessing.

## Declarations

### Ethical approval

Not applicable.

### Consent to participate

Not applicable.

### Consent for publication

Not applicable.

## Appendix A Supplementary Tables

**Table S1.** Some descriptive statistics and frequency observations of NRTI and NNRTI genotype-phenotype data from the Stanford HIV Drug Resistance Database.

| Drugs | Isolates | Muts | Max | Min | Mean | Std | Mode | Q1 | Q3 |
| --- | --- | --- | --- | --- | --- | --- | --- | --- | --- |
| 3TC | 2122 | 1336 | 100 | 0.500 | 47.587 | 46.257 | 100 | 1.500 | 100 |
| ABC | 2057 | 1270 | 100 | 0.500 | 4.640 | 6.036 | 0.900 | 1.200 | 6.100 |
| AZT | 2143 | 1348 | 100 | 0.100 | 21.629 | 35.102 | 100 | 0.700 | 24 |
| D4T | 2153 | 1350 | 100 | 0.400 | 2.297 | 4.485 | 0.800 | 0.900 | 2.400 |
| DDI | 2153 | 1350 | 100 | 0.400 | 2.049 | 3.810 | 1 | 1.100 | 2.100 |
| TDF | 1814 | 1262 | 100 | 0.200 | 1.807 | 3.945 | 0.800 | 0.800 | 1.700 |
| EFV | 2212 | 1361 | 100 | 0.100 | 23.807 | 37.074 | 100 | 0.600 | 30 |
| NVP | 2210 | 1357 | 100 | 0.100 | 40.323 | 45.284 | 100 | 0.700 | 100 |
| ETR | 984 | 1183 | 100 | 0.100 | 6.520 | 17.448 | 0.900 | 0.600 | 3.100 |
| RPV | 188 | 1159 | 4.605 | -1.609 | 1.064 | 1.585 | 4.605 | -0.053 | 2.028 |

**Table S2.** Performance metrics of the IF, DIF, and DIF-C models using 10-fold cross-validation methodology. The table displays the mean values of MSE, and *r* metrics across ten distinct test sets that are not used during the training phase.

| Classes | Drugs | Mean MSE | | | Mean $r$ | | |
| --- | --- | --- | --- | --- | --- | --- | --- |
|  |  | IF | DIF | DIF-C | IF | DIF | DIF-C |
| NRTIs | 3TC | 0.098 | 0.116 | 0.162 | 0.933 | 0.935 | 0.931 |
|  | ABC | 0.034 | 0.037 | 0.044 | 0.890 | 0.889 | 0.874 |
|  | AZT | 0.174 | 0.164 | 0.213 | 0.882 | 0.889 | 0.891 |
|  | D4T | 0.028 | 0.028 | 0.035 | 0.848 | 0.876 | 0.842 |
|  | DDI | 0.019 | 0.025 | 0.028 | 0.818 | 0.722 | 0.689 |
|  | TDF | 0.043 | 0.044 | 0.056 | 0.766 | 0.733 | 0.712 |
| NNRTIs | EFV | 0.184 | 0.200 | 0.195 | 0.888 | 0.854 | 0.891 |
|  | NVP | 0.517 | 0.237 | 0.242 | 0.653 | 0.864 | 0.873 |
|  | ETR | 0.279 | 0.226 | 0.256 | 0.861 | 0.696 | 0.677 |
|  | RPV | 0.373 | 0.229 | 0.232 | 0.544 | 0.800 | 0.833 |
| Mean NRTIs |  | 0.066 | 0.069 | 0.089 | 0.856 | 0.843 | 0.823 |
| Mean NNRTIs |  | 0.338 | 0.223 | 0.231 | 0.737 | 0.804 | 0.819 |
| All |  | 0.175 | 0.131 | 0.146 | 0.808 | 0.827 | 0.821 |

**Table S3.** Performance metrics of existing DIF-C models to capture the binary relationships between RTI pairs.

| ARVs |  | 3TC | ABC | AZT | D4T | DDI | TDF | EFV | NVP | ETR |
| --- | --- | --- | --- | --- | --- | --- | --- | --- | --- | --- |
| ABC | Accuracy | 0.984 |  |  |  |  |  |  |  |  |
|  | Sensitivity | 0 |  |  |  |  |  |  |  |  |
|  | Specificity | 1 |  |  |  |  |  |  |  |  |
|  | Precision | ~ |  |  |  |  |  |  |  |  |
|  | Recall | 0 |  |  |  |  |  |  |  |  |
|  | F1 Score | ~ |  |  |  |  |  |  |  |  |
| AZT | Accuracy | 0.904 | 0.834 |  |  |  |  |  |  |  |
|  | Sensitivity | 0.653 | 0.942 |  |  |  |  |  |  |  |
|  | Specificity | 0.991 | 0.704 |  |  |  |  |  |  |  |
|  | Precision | 0.959 | 0.794 |  |  |  |  |  |  |  |
|  | Recall | 0.653 | 0.942 |  |  |  |  |  |  |  |
|  | F1 Score | 0.777 | 0.861 |  |  |  |  |  |  |  |
| D4T | Accuracy | 0.990 | 0.996 | 0.863 |  |  |  |  |  |  |
|  | Sensitivity | 0 | 0.250 | 0.322 |  |  |  |  |  |  |
|  | Specificity | 1 | 0.999 | 0.996 |  |  |  |  |  |  |
|  | Precision | ~ | 0.500 | 0.951 |  |  |  |  |  |  |
|  | Recall | 0 | 0.250 | 0.322 |  |  |  |  |  |  |
|  | F1 Score | ~ | 0.333 | 0.481 |  |  |  |  |  |  |
| DDI | Accuracy | 0.989 | 0.994 | 0.868 | 0.897 |  |  |  |  |  |
|  | Sensitivity | 0 | 0.429 | 0.568 | 0.938 |  |  |  |  |  |
|  | Specificity | 1 | 0.998 | 0.997 | 0.858 |  |  |  |  |  |
|  | Precision | ~ | 0.600 | 0.987 | 0.862 |  |  |  |  |  |
|  | Recall | 0 | 0.429 | 0.568 | 0.938 |  |  |  |  |  |
|  | F1 Score | ~ | 0.500 | 0.721 | 0.898 |  |  |  |  |  |
| TDF | Accuracy | 0.992 | 0.992 | 0.901 | 0.884 | 0.834 |  |  |  |  |
|  | Sensitivity | 0 | 0 | 0.150 | 0.235 | 0.444 |  |  |  |  |
|  | Specificity | 1 | 1 | 0.998 | 0.957 | 0.963 |  |  |  |  |
|  | Precision | ~ | ~ | 0.900 | 0.381 | 0.798 |  |  |  |  |
|  | Recall | 0 | 0 | 0.150 | 0.235 | 0.444 |  |  |  |  |
|  | F1 Score | ~ | ~ | 0.257 | 0.291 | 0.570 |  |  |  |  |
| EFV | Accuracy | 0.930 | 0.931 | 0.884 | 0.937 | 0.947 | 0.959 |  |  |  |
|  | Sensitivity | 0.868 | 0.948 | 0.899 | 0.976 | 0.988 | 0.988 |  |  |  |
|  | Specificity | 0.955 | 0.911 | 0.870 | 0.862 | 0.879 | 0.875 |  |  |  |
|  | Precision | 0.886 | 0.923 | 0.875 | 0.930 | 0.932 | 0.957 |  |  |  |
|  | Recall | 0.868 | 0.948 | 0.899 | 0.976 | 0.988 | 0.988 |  |  |  |
|  | F1 Score | 0.877 | 0.935 | 0.887 | 0.953 | 0.959 | 0.973 |  |  |  |
| NVP | Accuracy | 0.910 | 0.928 | 0.913 | 0.920 | 0.916 | 0.919 | 0.784 |  |  |
|  | Sensitivity | 0.873 | 0.967 | 0.927 | 0.992 | 0.993 | 0.994 | 0.857 |  |  |
|  | Specificity | 0.935 | 0.863 | 0.889 | 0.741 | 0.739 | 0.640 | 0.092 |  |  |
|  | Precision | 0.898 | 0.920 | 0.931 | 0.905 | 0.898 | 0.911 | 0.900 |  |  |
|  | Recall | 0.873 | 0.967 | 0.928 | 0.992 | 0.993 | 0.994 | 0.857 |  |  |
|  | F1 Score | 0.885 | 0.943 | 0.929 | 0.947 | 0.943 | 0.951 | 0.878 |  |  |
| ETR | Accuracy | 0.897 | 0.844 | 0.848 | 0.826 | 0.830 | 0.864 | 0.866 | 0.939 |  |
|  | Sensitivity | 0.376 | 0.645 | 0.672 | 0.781 | 0.760 | 0.948 | 0 | 0.111 |  |
|  | Specificity | 0.990 | 0.930 | 0.965 | 0.867 | 0.879 | 0.764 | 0.973 | 0.991 |  |
|  | Precision | 0.875 | 0.799 | 0.929 | 0.845 | 0.814 | 0.827 | 0 | 0.444 |  |
|  | Recall | 0.376 | 0.645 | 0.672 | 0.781 | 0.760 | 0.948 | 0 | 0.111 |  |
|  | F1 Score | 0.526 | 0.714 | 0.780 | 0.811 | 0.786 | 0.884 | ~ | 0.178 |  |
| RPV | Accuracy | 0.802 | 0.767 | 0.868 | 0.833 | 0.819 | 0.911 | 0.789 | 0.937 | 0.571 |
|  | Sensitivity | 0.600 | 0.815 | 0.853 | 0.905 | 0.967 | 0.974 | 0.067 | 0.500 | 1 |
|  | Specificity | 0.873 | 0.694 | 0.903 | 0.500 | 0.409 | 0.500 | 0.982 | 1 | 0 |
|  | Precision | 0.625 | 0.800 | 0.955 | 0.893 | 0.819 | 0.927 | 0.500 | 1 | 0.571 |
|  | Recall | 0.600 | 0.815 | 0.853 | 0.905 | 0.967 | 0.974 | 0.067 | 0.500 | 1 |
|  | F1 Score | 0.612 | 0.807 | 0.901 | 0.899 | 0.887 | 0.950 | 0.118 | 0.667 | 0.727 |

**Table S4.**
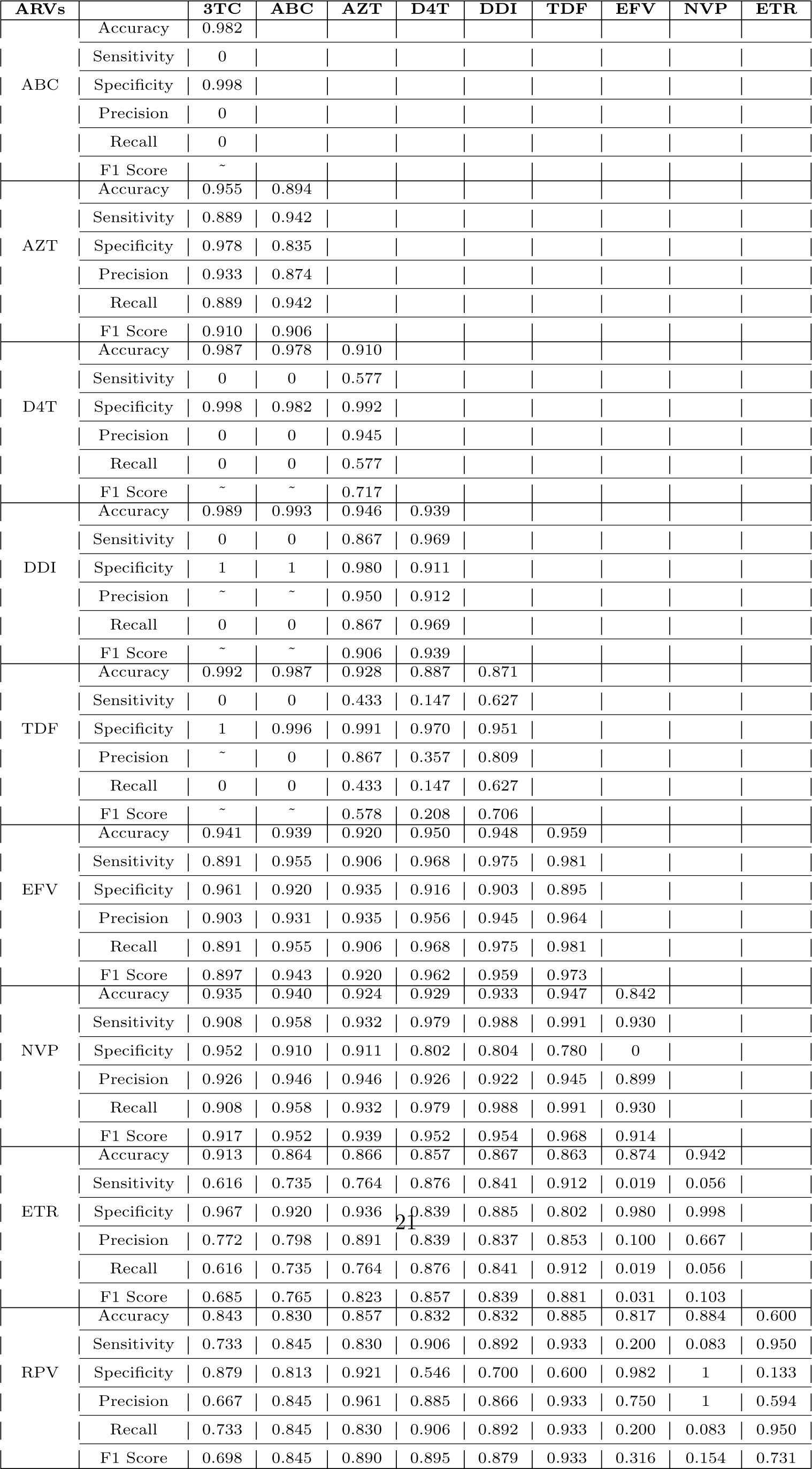
Performance metrics of existing DIF models to capture the binary relationships between RTI pairs.

## Appendix B Supplementary Figures

**Fig. S1.**
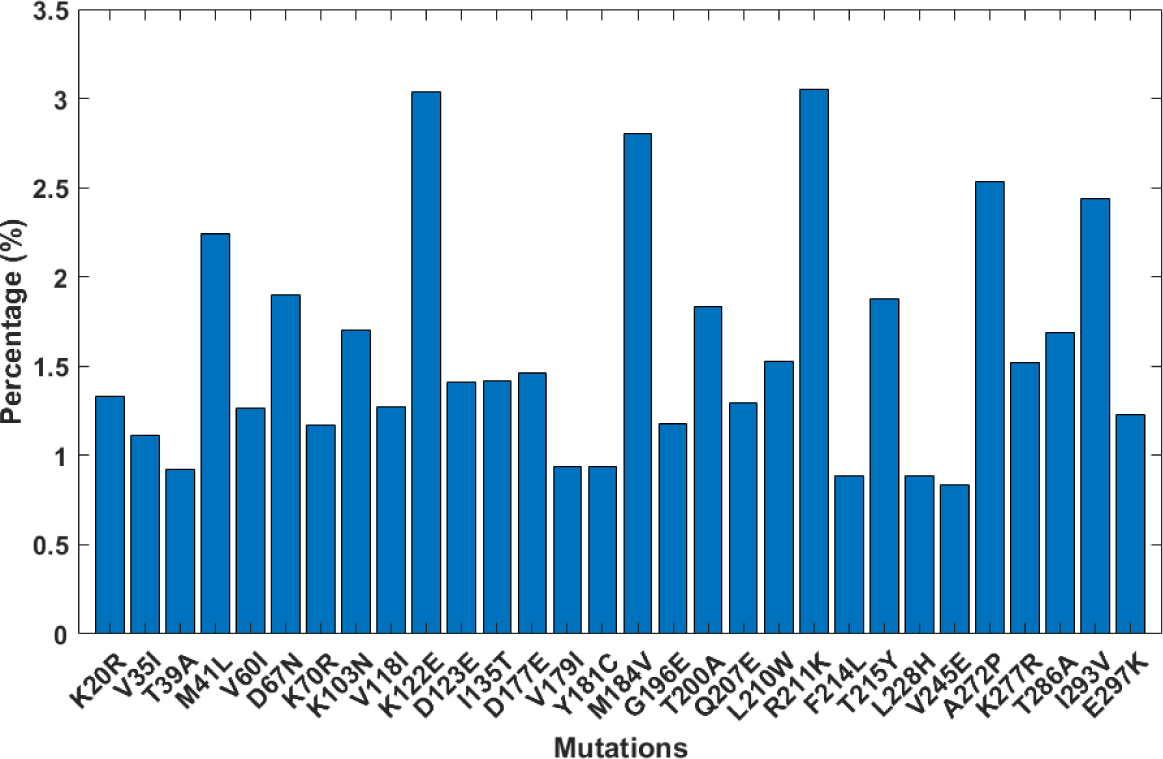
The top 30 RT mutations with the highest frequency among 1388 mutations identified, based on data from the Stanford HIV Drug Resistance Database.

**Fig. S2.**
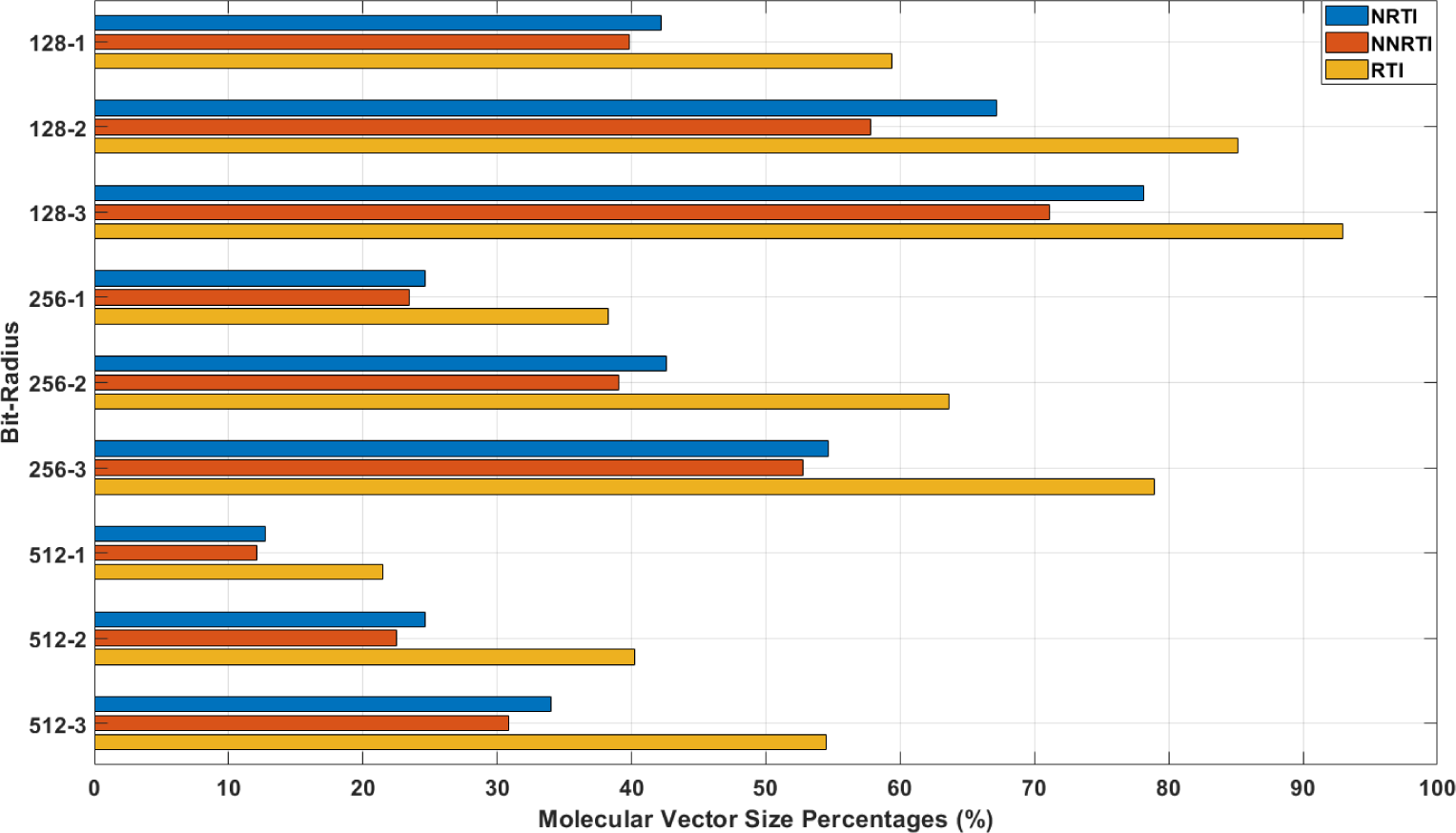
The significant vector sizes for different molecular representations are expressed as percentages. During the analysis, it was determined that within the 2-radius and 512-bit molecular representation, 126 out of the 512 bits hold unique properties for six NRTIs, 115 bits for four NNRTIs, and 206 bits for ten RTIs.

**Fig. S3.**
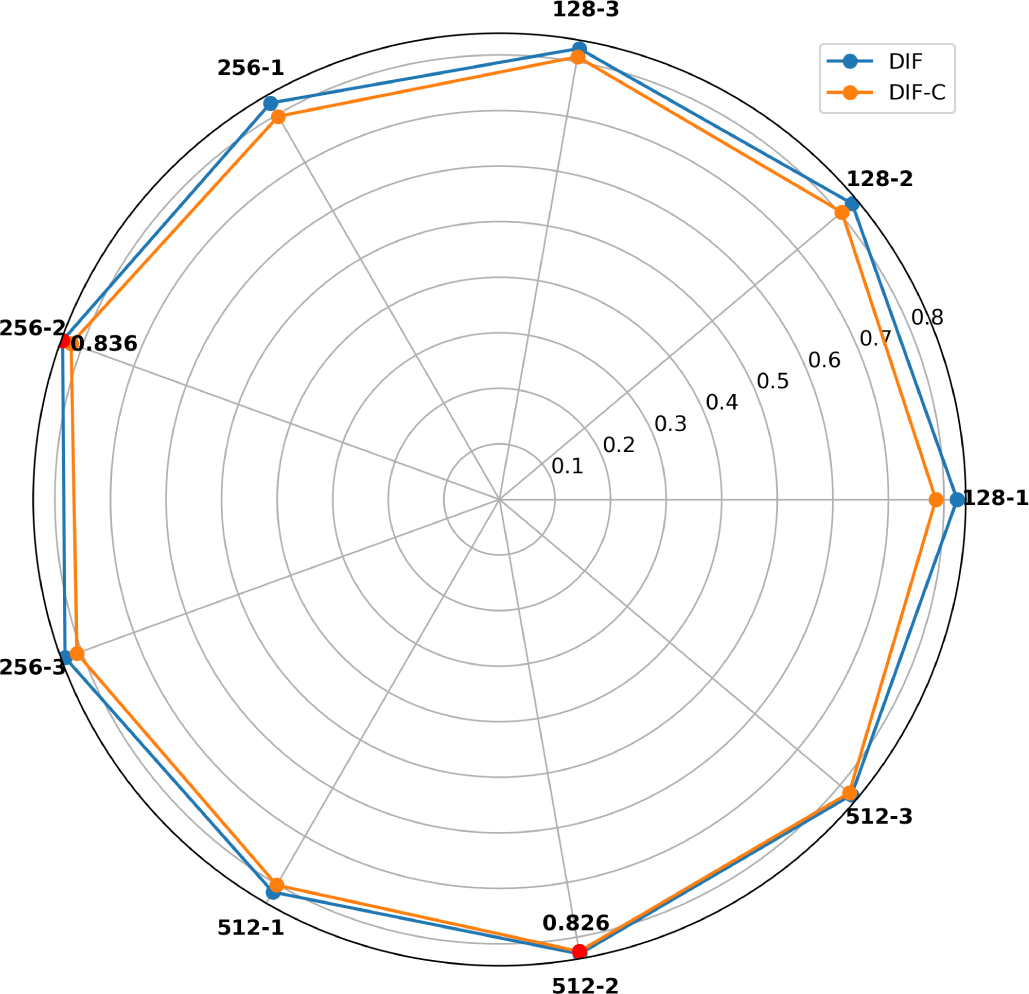
The optimal molecular representation for inhibitor modeling is determined by comparing vector sizes (128-bit, 256-bit, 512-bit) and radius values (1, 2, 3) across ten inhibitors in DIF and DIF-C models, as illustrated by the obtained *r* values. The red dots represent the highest *r* values obtained.

**Fig. S4.**
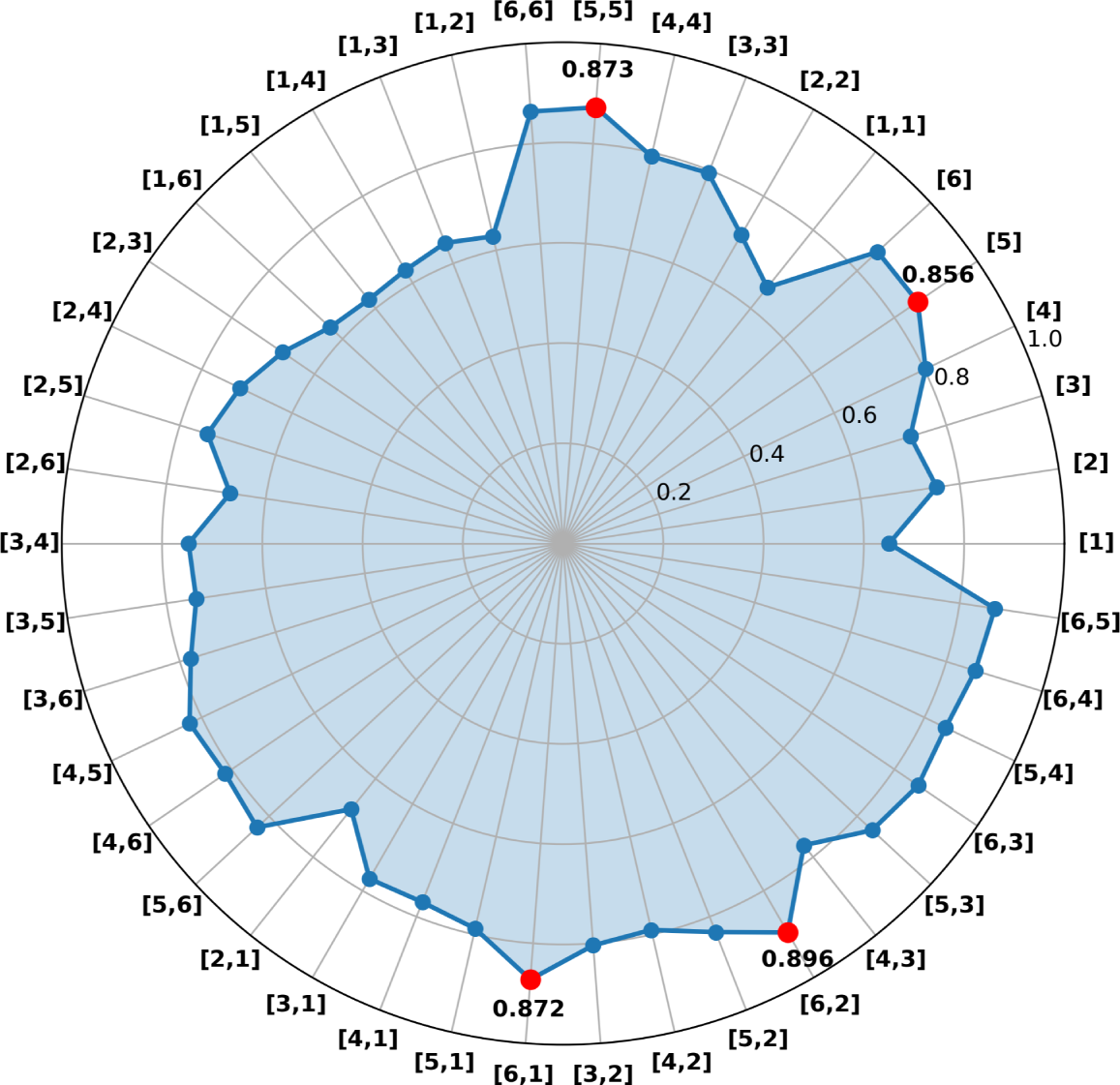
The illustrates mean *r* values for DIF models using various ANN architectures, where red dots represent mean *r* values for specific configurations. Evaluating 42 unique combinations by adjusting hidden layers (from 1 to 2) and neurons (from 1 to 6), the model with one hidden layer and 5 neurons is selected as optimal, balancing performance with computational efficiency.

**Fig. S5.**
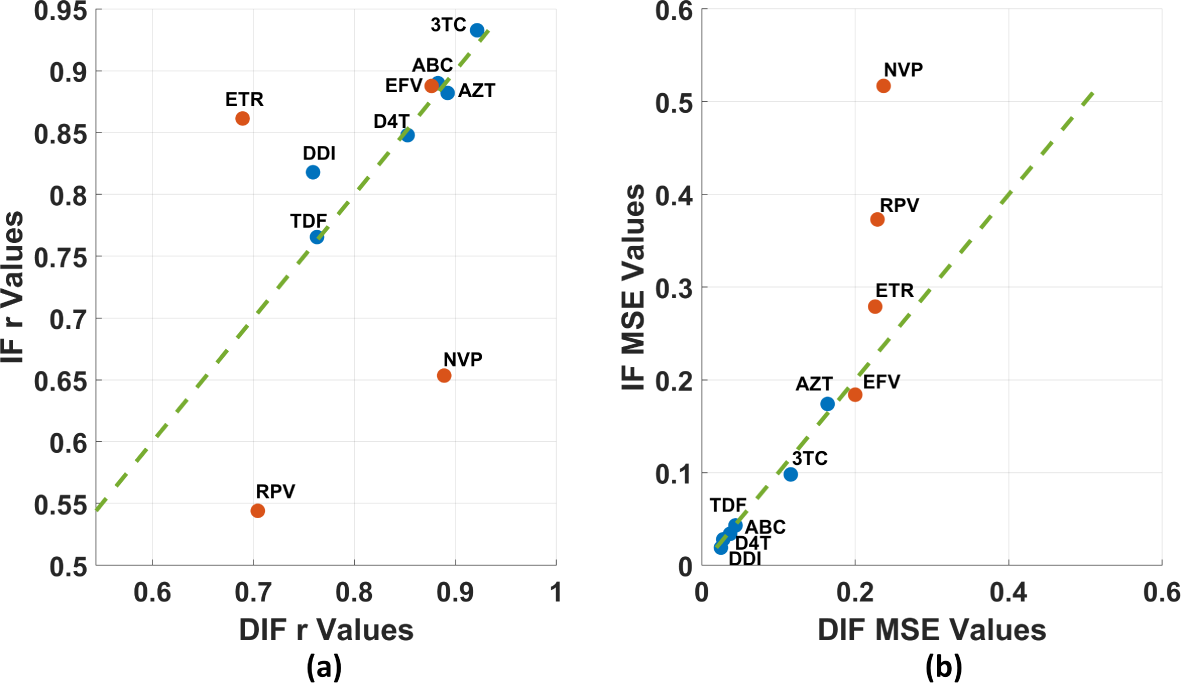
Comparative illustrations of (a) *r* and (b) MSE values for IF and DIF models. Both figures display the values of DIF models along the x-axis and the values of IF models along the y-axis. The blue markers represent the NRTI class, while the orange markers represent the NNRTI class.

**Fig. S6.**
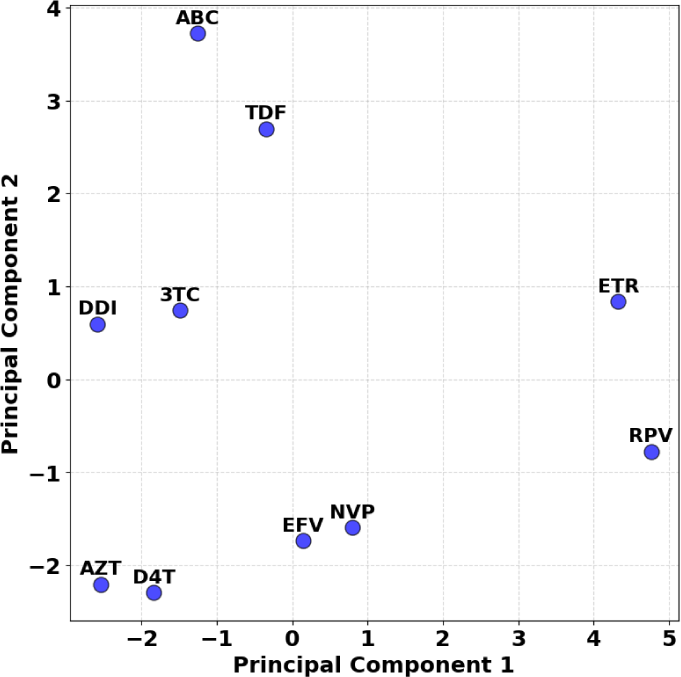
2D visualization of ten RTI molecules using 512-bit and 2-radius Morgan fingerprint representations by utilizing the PCA.

**Fig. S7.**
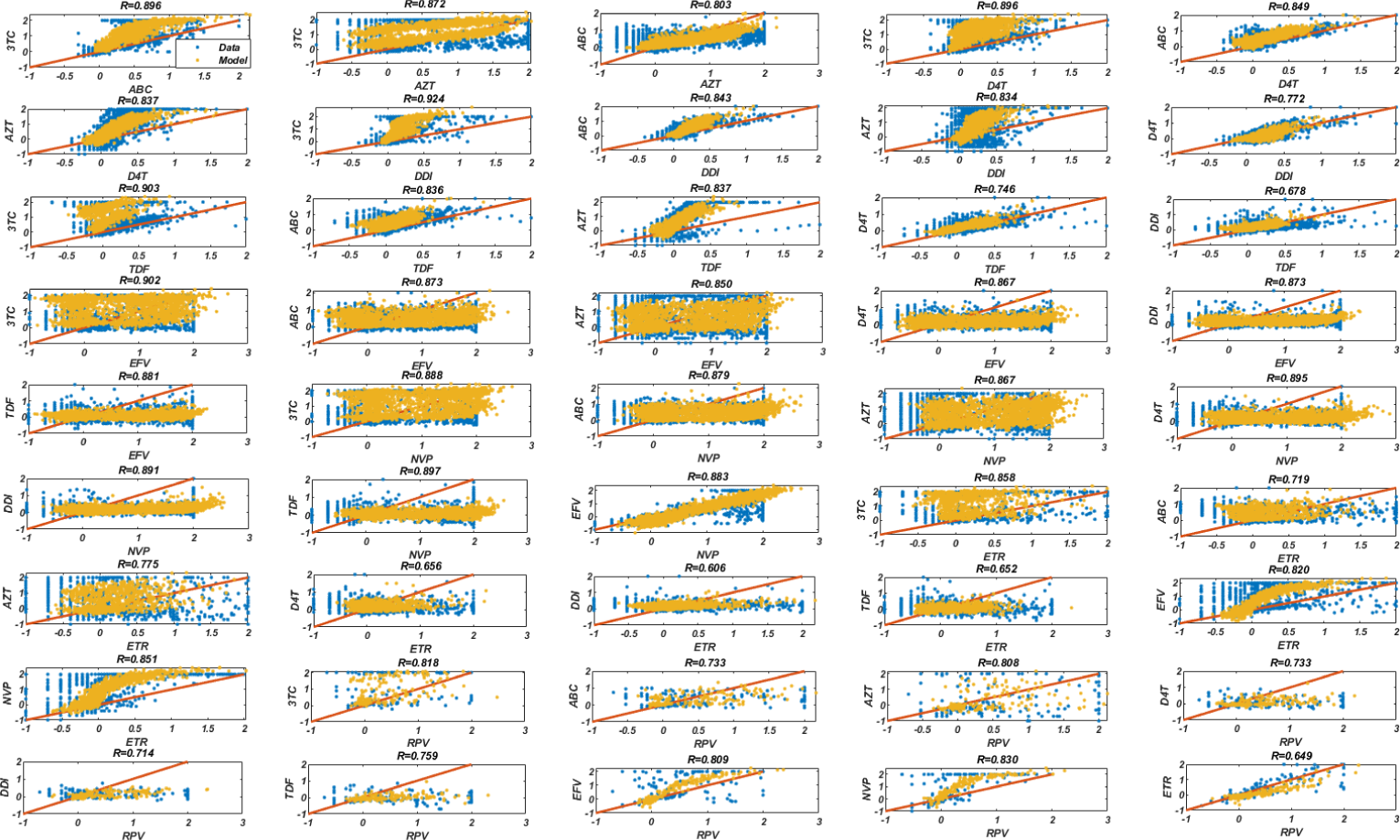
Prediction of fold change tendencies of inhibitor pairs with the DIF-C model obtained by training NRTIs and NNRTIs together.

